# Colorectum and Embedded Networks of Nerve Fibers Present Auxetic Responses During Uniaxial Circumferential Extension

**DOI:** 10.1101/2025.04.02.646859

**Authors:** Amirhossein Shokrani, Atta Seck, Kazunori Hoshino, Bin Feng, David M. Pierce

## Abstract

Understanding the multiscale mechanics of the colorectum is essential for uncovering the mechanotransductive pathways underlying visceral nociception. Intraluminal distension of the large intestine reliably evokes pain in disorders of gut-brain interaction (DGBIs), yet the tissue-level and nerve fiber-level responses to mechanical loading remain poorly defined. Here, we present results from a novel biomechanical testing framework that integrates uniaxial circumferential extension with high-resolution optical imaging to quantify deformation in both bulk colorectal tissue and embedded sensory nerve fibers. We tested intact, cylindrical colorectal segments from mice using a custom 3-D-printed chamber with intraluminal stainless-steel rods to apply circumferential stretch while maintaining a planar imaging field. We measured bulk-tissue deformation via Digital Image Correlation (DIC), while we assessed stretch in nerve fibers through fluorescence imaging of VGLUT2-labeled afferents analyzed using a custom fiber-network analyses. Across specimens, we observed a consistent auxetic response–characterized by positive axial strain during circumferential extension–at both the macroscale and microscale. Five out of six colorectal specimens exhibited positive axial Green-Lagrange strain (*E*_*xx*_), with an average median *E*_*xx*_ of 0.0177, during circumferential extension generating an average median *E*_*yy*_ of 0.1273. Nerve fiber analysis across nine specimens revealed an average median stretch ratio of 1.0631, indicating 6.31% elongation, with substantial heterogeneity driven by fiber orientation. These findings demonstrate that the colorectum and its embedded network of nerve fibers exhibit auxetic behavior, a property that may amplify mechanical signaling and influence nociceptive signaling. Our methods and results provide foundational insight into structure-function relationships of colorectum and inform design of bioinspired auxetic materials.

## 1. Introduction

The large intestine, consisting of the colon, rectum, and anus, functions as the terminal site of the gastrointestinal tract, where it absorbs water and electrolytes while forming solid feces from undigested residues. The colon primarily serves as a reservoir, regulating defecation to maintain homeostasis [1]. Additionally, the large intestine hosts a diverse gut microbiota that ferment undigested carbohydrates, producing short-chain fatty acids (SCFAs) such as acetate, propionate, and butyrate. These metabolites provide energy for colonic cells and support gut health [2, 3]. Microbial fermentation also generates gases that contribute to intraluminal distension, a key factor in abdominal and visceral pain [4, 5].

Intraluminal distension of the large intestine, particularly in the distal colorectum, reliably induces visceral pain in healthy individuals and exacerbates pain in patients with disorders of gut-brain interactions (DGBIs) [6, 7, 8]. Notably, stimuli such as cutting, pinching, and heating, which elicit significant pain in the skin, do not produce comparable nociceptive responses when applied to the small or large intestine. Since intraluminal distension serves as an adequate nociceptive stimulus in the colorectum, mechanotransduction by sensory afferents within the colon and rectum plays a critical role in visceral nociception [9, 10]. A deeper understanding of colorectal mechanotransduction during tissue deformation could facilitate the development of targeted therapies for DGBI-associated visceral pain [11, 12].

Colorectal mechanotransduction and perception of visceral pain involve multiscale interactions, where macroscale colorectal deformation induces microscale stress and strain in afferent fibers. At the macro scale, most biological tissues exhibit a positive Poisson’s ratio, contracting orthogonally to an applied uniaxial extension [13, 14, 15]. However, certain biological tissues display a negative Poisson’s ratio under mechanical extension, a phenomenon known as auxetic behavior, which results in lateral expansion orthogonal to the direction of distension. This auxetic response, deviating from typical soft tissue mechanics, has been observed in specific conditions within tendons and skin [16, 17].

At the micro scale, recent anatomical studies employing optical tissue clearing have precisely quantified afferent fiber density and morphology across distinct colorectum sublayers, identifying a high concentration of afferent fibers within the myenteric plexus and submucosa [18, 19, 20, 21]. Approximately 22% of colorectal afferents terminate within the ganglia of the myenteric plexus, a key component of the intrinsic nervous system in the gastrointestinal (GI) tract. These afferent endings within the myenteric plexus likely contribute to visceral nociception, highlighting their potential role in colorectal sensory processing [22, 23].

Nociceptors, a specialized subset of afferents, mediate pain perception by detecting injurious stimuli. Stretch-sensitive colorectal afferents, classified as nociceptors, encode colorectal distension through unmyelinated C-fibers that respond to noxious extension and exhibit sensitization [24, 10]. Their role in pain signaling is crucial to the pathophysiology of disorders such as irritable bowel syndrome (IBS), a disorder of gut-brain interaction (DGBI), where colorectal mechanical distension drives pain perception.

Understanding how mechanical forces influence colorectal nociception requires precise characterization of tissue and fiber deformations at multiple scales. Colorectal distension, and the resulting macroscale tissue deformations, generates complex micro-mechanical stimuli (micro-scale stresses and strains) affecting afferent fibers, yet the relationship between these deformations remains poorly defined. To address this gap, we established a novel framework for biomechanical testing to measure axial stretch in intact, cylindrical colorectal segments (10 mm long) subjected to controlled circumferential distension. We also fluorescently labeled extrinsic sensory afferents using a vesicular glutamate transporter type 2 (VGLUT2) promoter to drive tdTomato expression in transgenic mice, enabling optical tracking of sensory fiber deformation within the myenteric nervous system during controlled circumferential distension. We aimed to establish foundational datasets to enable theoretical models of colorectal tissue mechanics and to provide critical insights into how mechanical cues modulate nociceptive signaling, particularly in disorders such as irritable bowel syndrome (IBS). We hypothesize that macroscale colorectal deformation correlates with microscale afferent fiber strain by measuring and comparing both. The former employing tools for Digital Image Correlation (DIC) and the later employing our established framework for quantifying stretches within networks of neural fibers [25, 26]. Validating this relationship will enhance our understanding of nociceptor sensitization and inform the development of targeted therapies for visceral pain [27, 28, 29].

### 2. Materials and Methods

The University of Connecticut Institutional Animal Care and Use Committee reviewed and approved all experimental procedures, ensuring compliance with ethical standards for animal research. We housed mice in pathogen-free, American Association for Accreditation of Laboratory Animal Care (AAALAC)-accredited facilities, adhering to Public Health Service assurances and the Eighth Edition of the Guide for the Care and Use of Laboratory Animals. Individually ventilated polycarbonate cages (Animal Care System M.I.C.E.) accommodated up to five mice per cage, with environmental enrichment including nestlets and huts. Bedding material consisted of Envigo T7990 B.G. Irradiated Teklad Sani-Chips. Mice received ad libitum access to either 2918 Irradiated Teklad Global 18% Rodent Diet or 7904 Irradiated S2335 Mouse Breeder Diet (Envigo) and also received reverse osmosis water chlorinated to 2 ppm via water bottles. Animal care staff maintained housing conditions on a 12:12 light-dark cycle, with ambient temperature controlled between 70–77^*°*^F (set point: 73.5^*°*^F) and relative humidity regulated between 35–65% (set point: 50%). Animal care staff further conducted daily health checks, and changed cages biweekly to ensure optimal welfare.

### 2.1. Specimen preparation

### 2.1.1. Colorectal specimens for tracking deformation of bulk tissue

We used C57BL/6 wild-type mice of both sexes, aged 8—13 weeks and weighing 25— 30 g. We deeply anesthetized mice via isoflurane inhalation (2–5%) until the loss of the plantar reflex to forceps pinching confirmed adequate anesthesia. Once anesthetized, we placed the mice in a supine position and performed a midline laparotomy, initiating a 2 cm incision near the genitalia and extending proximally to expose the visceral cavity. We then incised the diaphragm to reveal the inferior view of the lungs and heart. We severed the right atrium and immediately immersed the mouse in chilled Krebs solution (in mM: 117.9 NaCl, 4.7 KCl, 25 NaHCO_3_, 1.3 NaH_2_PO_4_, 1.2 MgSO_4_, 2.5 CaCl_2_, and 11.1 D-glucose).

We then transferred the carcass to a dissection chamber containing fresh, chilled Krebs solution, where we harvested the distal 25 mm of the colorectum and carefully removed the mesentery and any residual connective tissue. For further structural analysis, we subdivided the colorectum into three distinct regions: colonic, intermediate, and rectal. We excised the rectal region (approximately 8 mm in length) while preserving the colonic and intermediate regions as continuous segments (approximately 17 mm long). We meticulously removed connective tissues and the mesenteric attachment to the colorectum.

We inserted two intraluminal stainless steel rods (25 mm and 30 mm in length, both with diameter of 1.2 mm) into the combined colonic and intermediate colorectum, as well as the rectal region, and inserted this assembly into a custom-built, 3-D-printed chamber with the mesenteric side facing towards the microscope objective. We pre-stretched the specimen circumferentially, by moving the shorter rod while the longer rod remained fixed in the chamber, just sufficient to ensure a planner surface for subsequent application of speckle pattern and imaging. We then used a gas ball valve to blow air onto the colon, drying it just until the surface appeared matte. To apply the speckle pattern we utilized a dual action Iwata airbrush and air compressor (Custom Micron Plus Gravity Feed Dual Action Airbrush and Studio Series Sprint Jet Single Piston Air Compressor, ANEST Iwata-Medea, Inc., Portland, OR). We mixed black powdered pigment (Pearl Ex Powder Pigments: 640 Carbon Black, Jacquard Products, Healdsburg, CA) with deionized (DI) water at a 2:5 ratio. The final solution had a total volume of approximately 3 ml which we poured into the gravity feed cup attached to the airbrush and mixed via pipetting. Subsequently, we placed the chamber with the slightly dried colorectum in an enclosure and sprayed the pigment on from *∼*15 cm until the colon was sparsely and evenly covered. We then dried the specimen again and cleaned the chamber using Kimwipes (Kimtech Science, Kimberly-Clark, Irving, TX). Once clean, we filled the chamber with Krebs solution.

#### 2.1.2. Colorectal specimens for tracking deformation of nerve fibers

We used transgenic mice heterozygous for the VGLUT2-Cre and tdTomato (VGLUT2/tdT) genes, of both sexes and aged 18–35 weeks and 20–30 g. We anesthetized mice using isoflurane inhalation (2–5%) until the absence of plantar reflex response to forceps pinching. Subsequently, we euthanized mice using transcardial perfusion with oxygenated Krebs solution (in mM: 117.9 NaCl, 4.7 KCl, 25 NaHCO_3_, 1.3 NaH_2_PO_4_, 1.2 MgSO_4_, 2.5 CaCl_2_, and 11.1 D-glucose at room temperature and bubbled with 95% O_2_ and 5% CO_2_) from the left ventricle to the right atrium. We carefully harvested the distal 22 mm from the anal ring of the large intestine while keeping the mouse submerged in fresh oxygenated Krebs solution. We then meticulously removed connective tissues and the mesenteric attachment to the colorectum. Finally, we inserted two intraluminal stainless steel rods (25 mm and 30 mm in length) into the combined colonic and intermediate colorectum, as well as the rectal region, and inserted this assembly into a custom-built, 3-D-printed chamber with the mesenteric side facing towards the microscope objective, and we filled the chamber with Krebs solution.

### 2.2. Mechanical testing with integrated imaging

We applied uniaxial, circumferential mechanical extension to the intact, tubular colorectum using two intraluminal cylindrical rods. The smooth stainless steel rods minimally influenced the axial load and deformation of the colorectum, allowing controlled circumferential stretch while maintaining the colorectal surface in-plane for optimal optical imaging (see fig. 1). We inserted the colorectum with intraluminal rods on a glass layer within the 3-D printed chamber to minimize friction. We then filled the chamber with oxygenated Krebs solution at room temperature to maintain the viability of the colorectum tissue and the embedded nerve fibers. One of the inserted rods remained fixed, while we used a servo-controlled force actuator (Model 305D, Aurora Scientific, Aurora, Canada) to pull the other rod.

**Figure 1:**
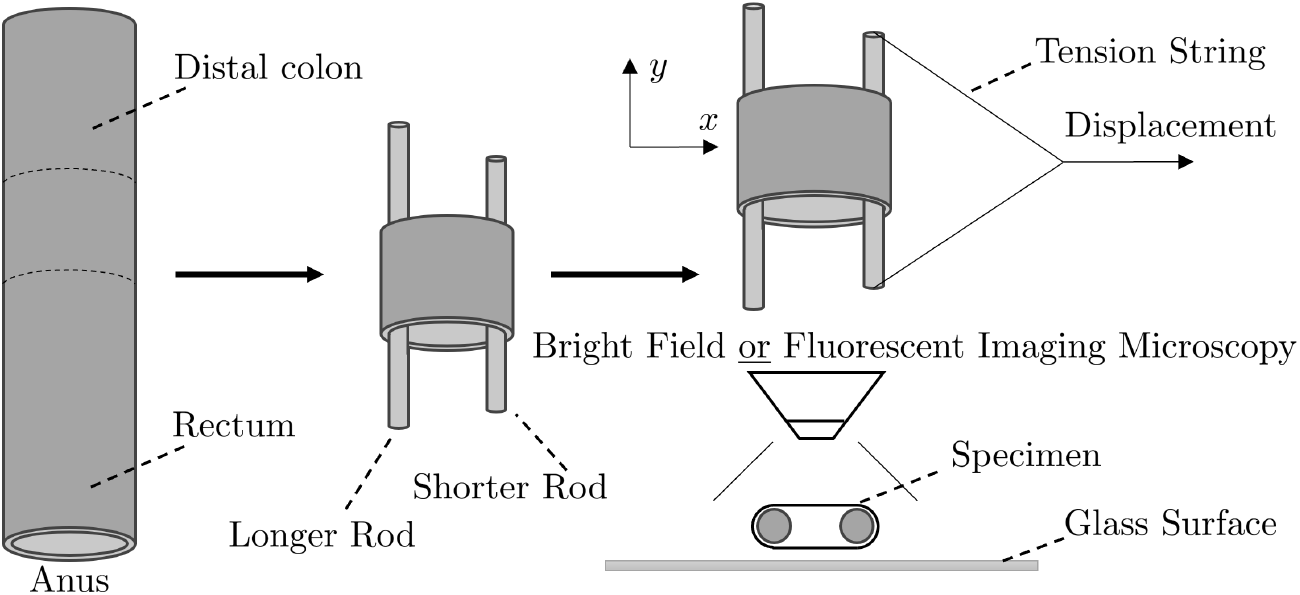
Schematic of the colorectal specimen and experimental setup for mechanical testing with integrated imaging. We imaged the deformation of the carbon-sprayed bulk tissue using bright-field microscopy (BF) and we imaged the deformation of the network of nerve fibers using fluorescent imaging microscopy (FI) during circumferential, colorectal extension. We used a custom-built, 3-D-printed chamber to distend the colorectum with two intraluminal stainless-steel rods measuring 25 mm and 30 mm in length. The longer rod remained fixed while we applied the stretch using the shorter rod. The entire assembly glides over glass surfaces on the bottom and sides of the chamber to minimize friction.

During mechanical testing we used an upright fluorescent microscope (BX51WI, Olympus, Inc., Tokyo, Japan) with a 4*×* objective lens (Model UPlanFLN *∞*/ - / FN26.5, NA 0.13, Olympus) for both bright-field and fluorescent imaging. To record the colorectum and networked nerve fibers during the quasi-static ramped stretching, we employed the open-source microscopy software Micro-Manager [30] and captured images with a low-noise sCMOS camera (Zyla-4.2P, 82% quantum efficiency, Oxford Instruments Andor Ltd., Belfast, Northern Ireland) at ten frames per second with a resolution of 2048*×*2048 pixels. The raw image stack of tiff file recordings captured a 5.35*×*5.35 mm field of view.

#### 2.2.1. Tracking deformation with bright-field imaging of carbon-dot speckles

We positioned the chamber for application of circumferential stretch under the objective, with the colorectum mounted and submerged in Krebs solution. We pre-stretched the colorectum using a force of 3.65 *±* 3.03 mN. Using the Spike 2 software application (Spike2 version 5.02, Cambridge Electronic Design, Cambridge, UK), we applied a quasistatic, displacement-controlled ramp that increased 0–2 mm over 15 seconds, then returned to 0 mm over the following 15 seconds. To ensure repeatable results, we pre-conditioned the specimen by repeating the ramped-stretch protocol 3–5 times [31, 32]. We observed no significant changes in the mechanical integrity of the colorectum during pre-conditioning or multiple trials on the same colorectal specimen.

We cut the dissected 25 mm segment of colorectum into two segments: one containing the colonic and intermediate regions, and the other containing only the rectal region. For the colonic and intermediate regions, after pre-conditioning, we first adjusted the objective to focus on the colonic region and recorded one quasi-static cycle of ramped stretch. We then refocused the objective on the intermediate region and repeated the cycle. Although the colorectal segment containing both the colonic and intermediate regions underwent more cycles than the rectal segment alone, no significant differences in mechanical integrity were observed between the two layers. Once we completed all the recordings for the mesenteric side, we removed the Krebs solution, rotated the colorectum to expose the anti-mesenteric side, and repeated the entire procedure for the anti-mesenteric side.

We used an upright fluorescent microscope in bright field mode to visualize and record the deformation of the colorectal segments during circumferential colorectal stretching.

#### 2.2.2. Tracking deformation with fluorescent imaging of nerve fibers

We positioned the chamber for application of circumferential stretch under the objective, with the colorectum mounted and submerged in Krebs solution. We pre-stretched the colorectum using a force of 188.99 *±* 38.30 mN and applied a quasi-static, displacement-controlled ramp from 0 to 2 mm in 15 seconds [32]. We pre-conditioned the specimen by repeating this ramped stretch 3–5 times before recording the force-displacement response [31, 32]. Throughout the entire procedure, we observed no significant changes in the mechanical integrity of the colorectum.

We used an upright fluorescent microscope to visualize and record the motion of VGLUT2-labeled unmyelinated nerve fibers during circumferential colorectal stretching. Imaging the colonic and rectal regions for both mesenteric and antimesenteric sides required 2—4 cycles of quasi-static, ramped stretching (again 0 to 2 mm in 15 sec).

### 2.3. Image analyses and corresponding strain measurements

#### 2.3.1. Digital Image Correlation of bulk tissue deformation

We established custom Digital Image Correlation (DIC) tools to analyze the bulk tissue deformation of the speckle-dotted colorectal specimens [33, 34]. Briefly, we conducted optical tracking for each node of the model from the initial (reference) image to the final image and calculated a 2-D strain map from the optical analysis. We defined a small panel centered around each node in the reference image. We then computed the normalized cross-correlation (NCC) for areas around each node in the subsequent image to identify the best-matched pattern and calculate a displacement vector. Despite pixelation of the images, we achieved subpixel resolution for displacement using second-degree polynomial interpolation along both directions in 2-D. From our displacement vectors we then calculated distributions of deformation gradients and finally distributions of Green-Lagrange strains.

We implemented the DIC algorithm within MATLAB (V2024, Mathworks, Natick, MA) to track the carbon-sprayed speckle patterns and subsequently calculate distributions of Green-Lagrange strains [34].

#### 2.3.2. Quantitative analyses of nerve fiber deformation

We established a semi-automated framework to quantify and analyze the micron-scale patterns of stretches within the network of nerve fibers within the myenteric plexus undergoing macroscopic circumferential stretch [32]. Briefly, we extract the fiber network from initial (reference) images of the colorectum as a point cloud with connectivities representing the nerve fibers. Our algorithm then automatically tracks the deformation of the fiber network using images recorded from subsequent time points during the deformation and determines the stretch ratio of individual fiber bundles comprising the network. Finally, our algorithm generates plots of the stretch ratios as spatial heat maps and facilitates some quantitative analyses, e.g., stretch ratios vs. fiber angles.

We implemented the framework within MATLAB (V2024, Mathworks) and provide free, public access, including input files for a simple test case, at github.uconn.edu/imLab/Fiber-Network_Analyses.

### 2.4. Statistical Analyses

#### 2.4.1. Digital Image Correlation of bulk tissue deformation

We conducted statistical analyses to quantify the mechanical response of colorectal specimens undergoing uniaxial circumferential extension. First, we discretized the central region of the imaged colorectal area (1.94*×*2.57 mm^2^) into 336 rectangular elements (16*×*21) and calculated the *E*_*xx*_ and *E*_*yy*_ in-plane, Green-Lagrange strains in each element from the nodal displacements throughout the entire mechanical test. We then determined the median and interquartile range (IQR) for each specimen to evaluate variability in the deformation response. Additionally, we computed the average *E*_*xx*_ and *E*_*yy*_ Green-Lagrange strains, and IQR, from the imaged area across all specimens.

#### 2.4.2. Quantitative analyses of nerve fiber deformation

We performed statistical analysis on the fiber-wise stretch ratios derived from our Neural Fiber Analysis Framework to assess the primary mechanical response of neural fibers during uniaxial circumferential extension. Using stretch values from the Fiber Network Analysis, we calculated the median stretches (*λ*) and the interquartile range (IQR) for each specimen to quantify variability in the fiber response under loading. Additionally, we determined the average stretches (*λ*), and IQR, from the imaged area across all specimens.

## 3. Results

We successfully completed two different biomechanical tests with integrated imaging. We tested six segments from the distal colon with carbon speckle-dots and analyzed the deformation of the bulk tissue using DIC. We also tested nine fluorescently labeled segments from the distal colon and analyzed the patterns of stretches within the network of nerve fibers. All specimens underwent uniaxial circumferential extension with median circumferential Green-Lagrange strains *E*_*yy*_ up to 0.18.

### 3.1. Digital Image Correlation of bulk tissue deformation

Five of the six colorectal specimens presented axial expansion with median axial Green-Lagrange strains *E*_*xx*_ up to 0.04, i.e., an auxetic response with negative Poisson’s ratio. The remaining specimen presented an insignificant axial expansion (*E*_*xx*_ *<* |0.005|), interpreted as a Poisson’s ratio of approximately zero.

In Fig. 2 we present images and analyses of a representative specimen from a mouse distal colon undergoing uniaxial circumferential extension, note the highlighted rectangular area used for DIC analysis (Fig. 2(a), 1.94 *×*2.57 mm^2^). In Fig. 2(b) we present the pattern of displacement vectors at each node used in the analyses, showing expansion in both circumferential and axial directions. In Figs. 2(c),(d) we provide the Green-Lagrange strains in the circumferential *E*_*yy*_ and axial *E*_*xx*_ directions, respectively. Positive axial strain is apparent throughout the area of the distal colon, indicating a negative Poisson’s ratio in response to circumferential extension.

**Figure 2:**
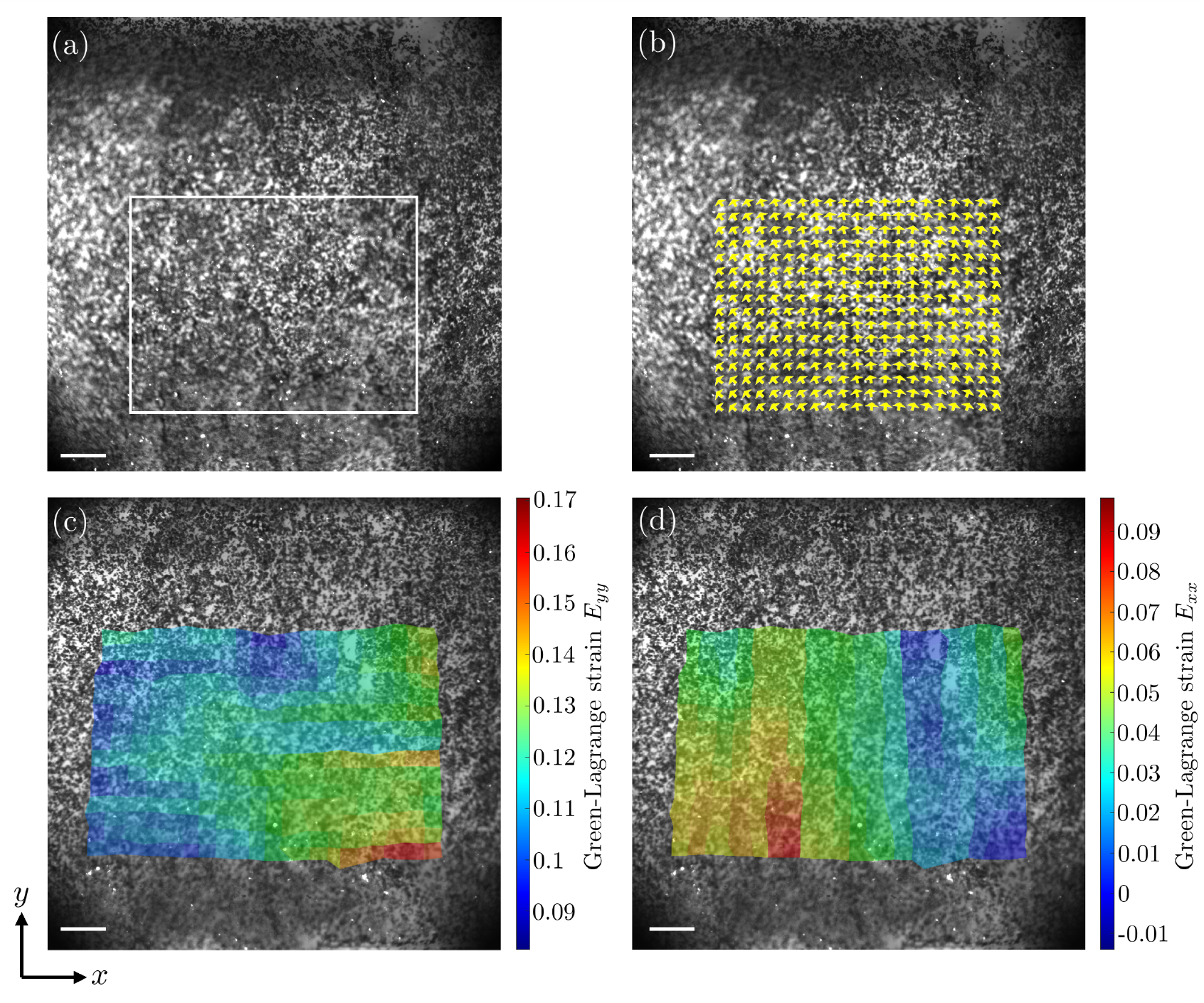
Auxetic mechanical response, i.e. negative Poisson’s ratio, of a representative specimen (#4 in Table B.1) from a mouse distal colon undergoing uniaxial circumferential extension in the *y*-direction. (a) Carbon speckle-dotted specimen in the reference configuration with the white box highlighting the area for Digital Image Correlation (DIC) analysis. (b) The distribution of displacement (unit) vectors for each node of the DIC analysis. (c) The distribution of Lagrange Green strains in the circumferential direction (*E*_*yy*_) calculated using DIC. (d) The distribution of Lagrange Green strains in the axial direction (*E*_*xx*_) calculated using DIC. The image field of view is 4.17*×*4.17 mm^2^ and the scale bar equals 417 *µ*m.

In Fig. 3 we present images and analyses of the one of six colorectal specimens that showed insignificant axial expansion (very near zero) in response to uniaxial circumferential extension, note the highlighted rectangular area used for DIC analysis (Fig. 3(a). In Fig. 3(b) we present the pattern of displacement vectors at each node used in the analyses, showing expansion in the circumferential but not axial directions. In Figs. 2(c),(d) we provide the Green-Lagrange strains in the circumferential *E*_*yy*_ and axial *E*_*xx*_ directions, respectively, indicating a Poisson’s ratio of approximately zero.

**Figure 3:**
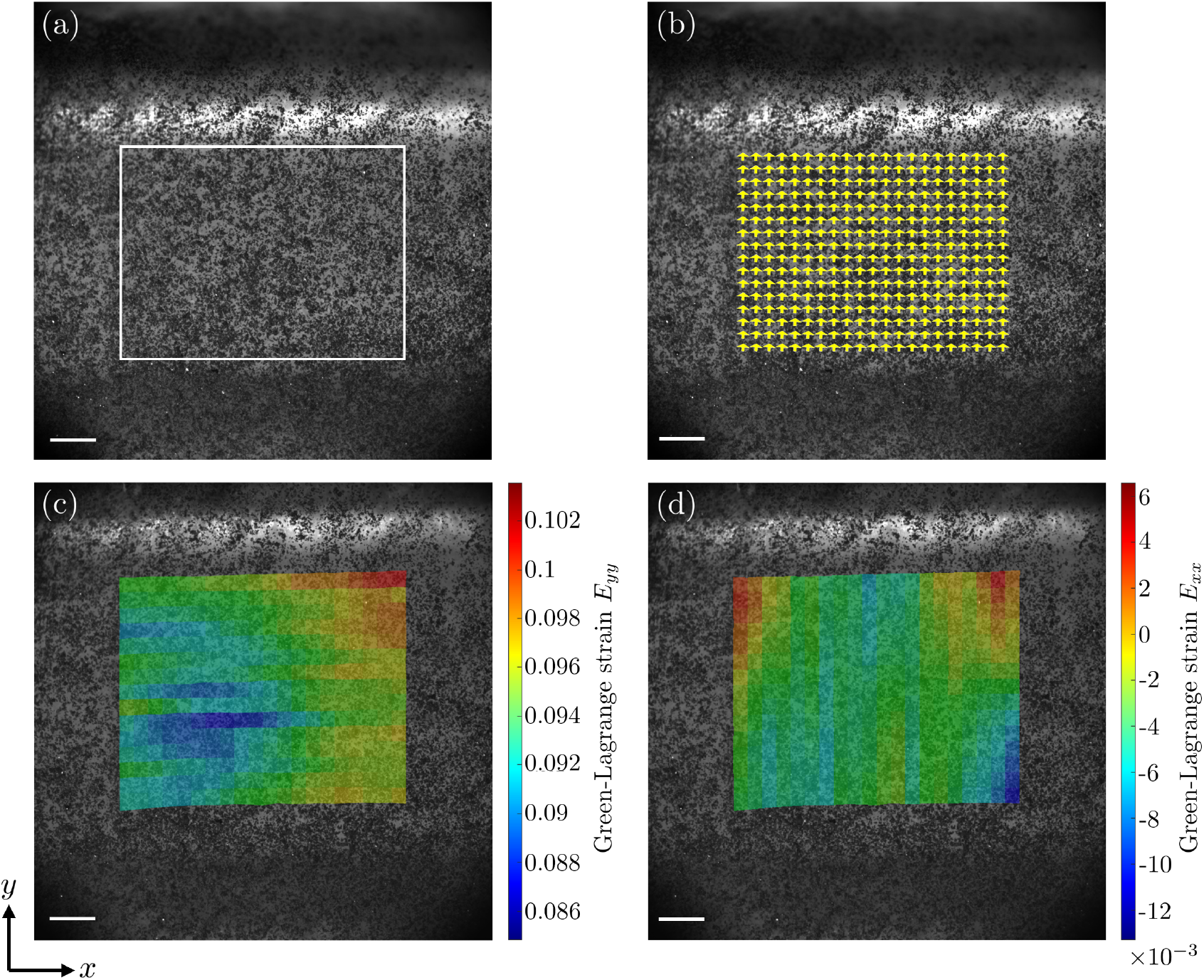
Mechanical response presenting Poisson’s ratio approximately zero of a representative specimen (#5 in Table B.1) from a mouse distal colon undergoing uniaxial circumferential extension in the *y*-direction. (a) Carbon speckle-dotted specimen in the reference configuration with the white box highlighting the area for Digital Image Correlation (DIC) analysis. (b) The distribution of displacement (unit) vectors for each node of the DIC analysis. (c) The distribution of Lagrange Green strains in the circumferential direction (*E*_*yy*_) calculated using DIC. (d) The distribution of Lagrange Green strains in the axial direction (*E*_*xx*_) calculated using DIC. The image field of view is 4.17 *×* 4.17 mm^2^ and the scale bar equals 417 *µ*m.

### 3.2. Quantitative analyses of nerve fiber deformation

In Fig. 4 we present a representative magnified image of fluorescent nerve fibers from a VGLUT2/tdT mouse colorectum, along with the overlaid map of fiber stretch ratios, showing expansion in both circumferential and axial directions during uniaxial circumferential extension. In Fig. 4(a) we present a detailed magnified view of VGLUT2-labeled unmyelinated nerve fibers from a VGLUT2/tdT mouse colorectum, covering a 3.57*×*3.57 mm^2^ field of view, with a box highlighting the area selected for further analyses. In Fig. 4(b) we present a heat map of the fiber stretch ratios during circumferential stretching of the colorectum. In Figs. 4(c),(d), we illustrate the stretch ratio for each fiber and analyze these ratios based on the fiber’s angle relative to the *x*-axis (axial direction within the colorectum). The specimen shown in Fig. 4 presented 7.25% lateral expansion of the fiber network embedded within the tissue during uniaxial circumferential extension of 2 mm.

**Figure 4:**
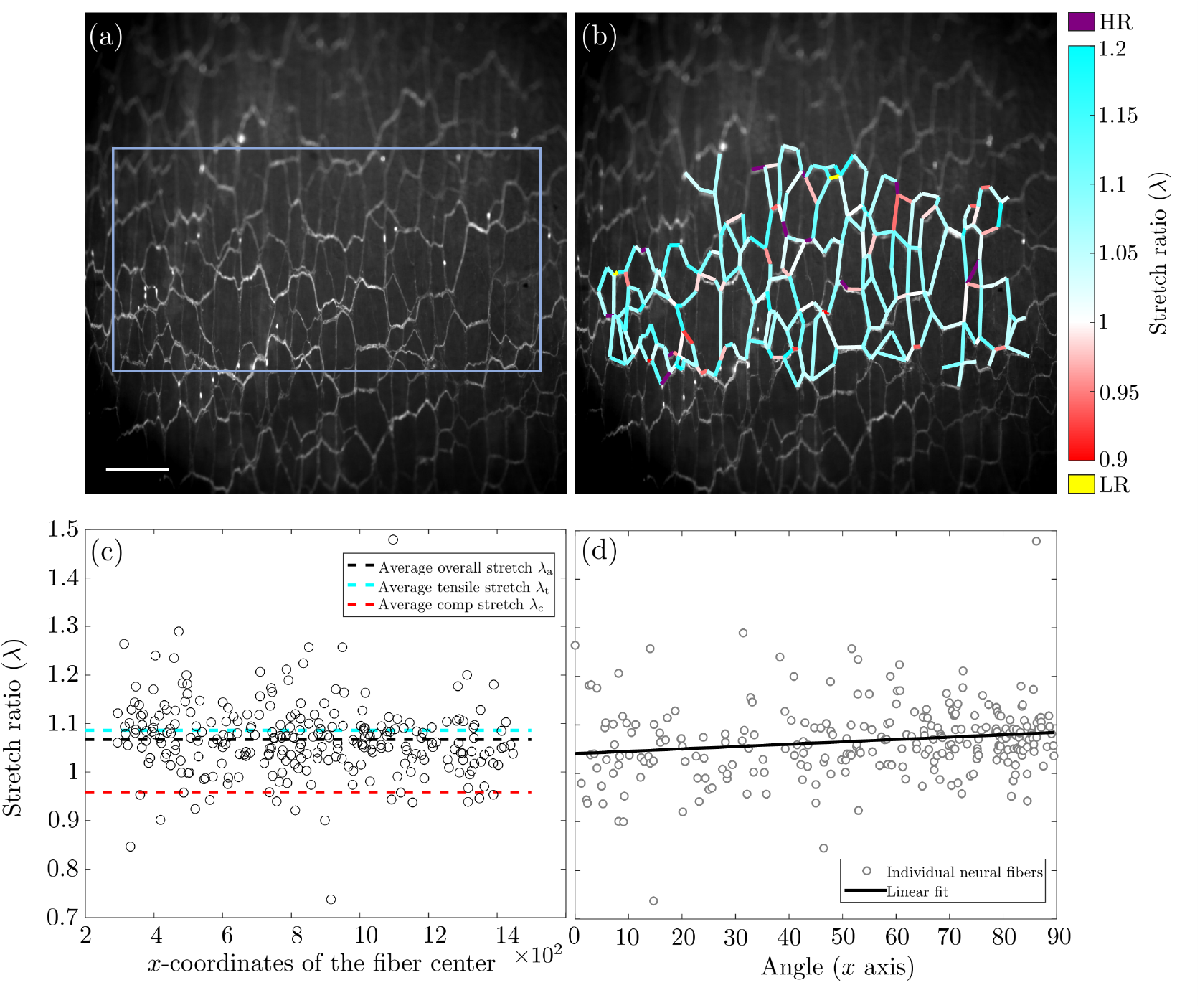
The distribution of stretch ratios of individual unmyelinated nerve fibers within a representative specimen (#XX in Table B.2) from a mouse distal colon undergoing uniaxial circumferential extension in the *y*-direction. This specimen presents an overall auxetic mechanical response, i.e. negative Poisson’s ratio. (a) VGLUT2-labeled nerve fibers in the reference configuration with light blue box highlighting area for subsequent analysis. (b) The distribution of fiber stretch ratios in the final configuration. HR: Higher Range (stretch ratios above 1.2); LR: Lower Range (stretch ratios below 0.9). (c) Stretch ratios for individual nerve fibers within the network. Black dashed line indicates average overall stretch ratio (*λ*_a_ = 1.067). Cyan and red dashed lines indicate the average tensile (*λ*_t_ = 1.086) and compressive (*λ*_c_ = 0.958) stretch ratios, respectively. (d) Linear correlation between stretch ratios and orientations of individual nerve fibers. An angle of 0^*°*^ indicates axial direction while 90^*°*^ indicates circumferential direction within the colorectum. The scale bar equals 465 *µ*m.

In Fig. 5 we present a magnified image of fluorescent nerve fibers from a representative mouse distal colon undergoing uniaxial circumferential extension, along with the overlaid map of fiber stretch ratios, showing expansion in the circumferential but not axial directions during uniaxial circumferential extension. In Fig. 5(a) we present a detailed magnified view of VGLUT2-labeled unmyelinated nerve fibers from a VGLUT2/tdT mouse colorectum, with a box highlighting the area selected for further analyses. In Fig. 5(b) we present a heat map of the fiber stretch ratios during circumferential stretching of the colorectum. In Figs. 5(c),(d), we illustrate the stretch ratio for each fiber and analyze these ratios based on the fiber’s angle relative to the *x*-axis (axial direction within the colorectum).

**Figure 5:**
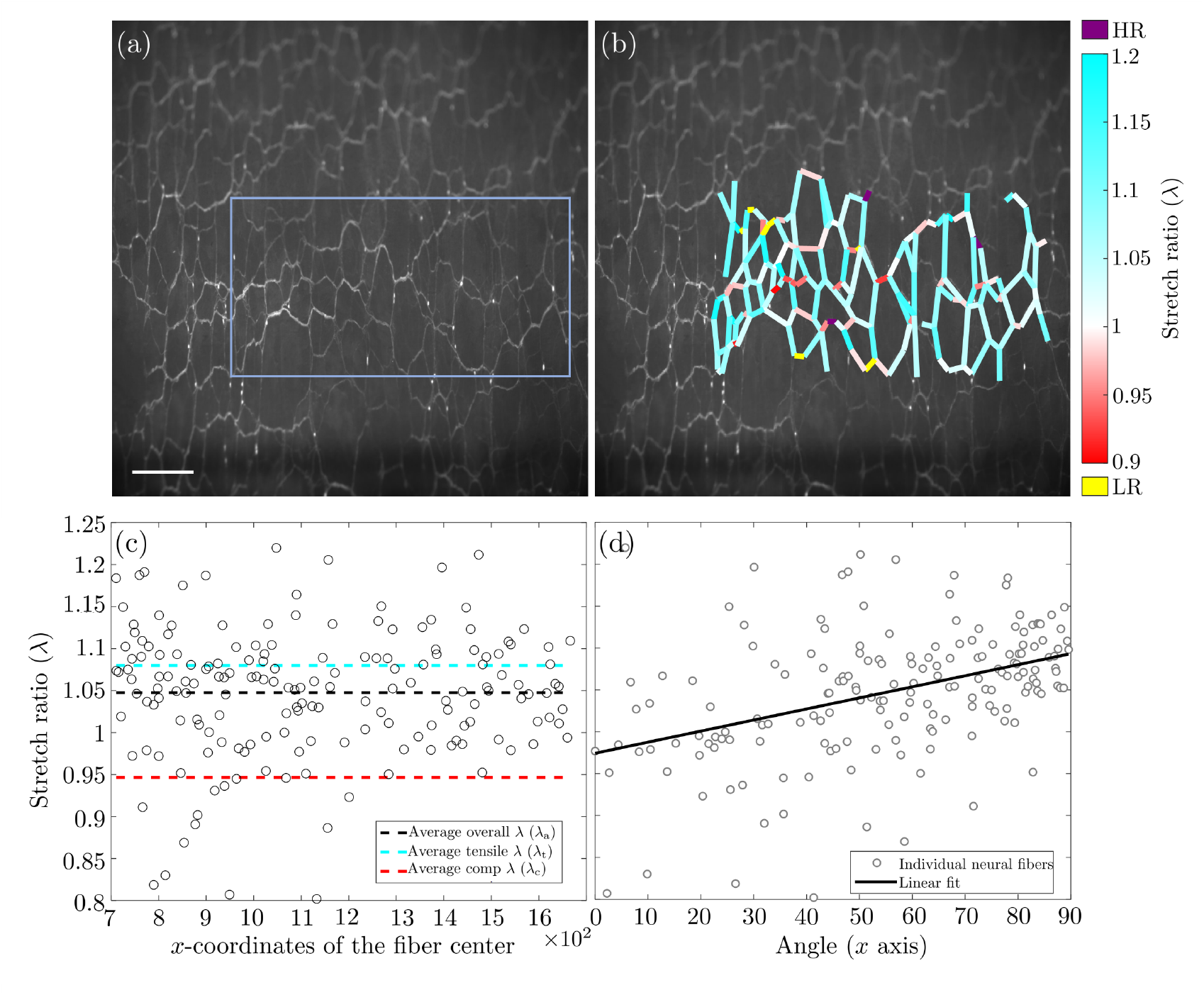
The distribution of stretch ratios of individual unmyelinated nerve fibers within a representative specimen (#XX in Table B.2) from a mouse distal colon undergoing uniaxial circumferential extension in the *y*-direction. This specimen presents an overall mechanical response showing a Poisson’s ratio of approximately zero. (a) VGLUT2-labeled nerve fibers in the reference configuration with light blue box highlighting area for subsequent analysis. (b) The distribution of fiber stretch ratios in the final configuration. HR: Higher Range (stretch ratios above 1.2); LR: Lower Range (stretch ratios below 0.9). (c) Stretch ratios for individual nerve fibers within the network. Black dashed line indicates average overall stretch ratio (*λ*_a_ = 1.048). Cyan and red dashed lines indicate the average tensile (*λ*_t_ = 1.080) and compressive (*λ*_c_ = 0.947) stretch ratios, respectively. (d) Linear correlation between stretch ratios and orientations of individual nerve fibers. An angle of 0^*°*^ indicates axial direction while 90^*°*^ indicates circumferential direction within the colorectum. The scale bar equals 465 *µ*m.

Additionally, we provide representative magnified fluorescent microscopy images of VGLUT2-labeled nerve fibers during circumferential extension of the colon in the reference and final configurations in Appendix A.

### 3.3. Statistical analyses

In Fig. 6 we present the median and interquartile range (IQR) of the Green-Lagrange strains in the circumferential *E*_*yy*_ and axial *E*_*xx*_ directions for all six colorectal specimens used for analyses via DIC.

**Figure 6:**
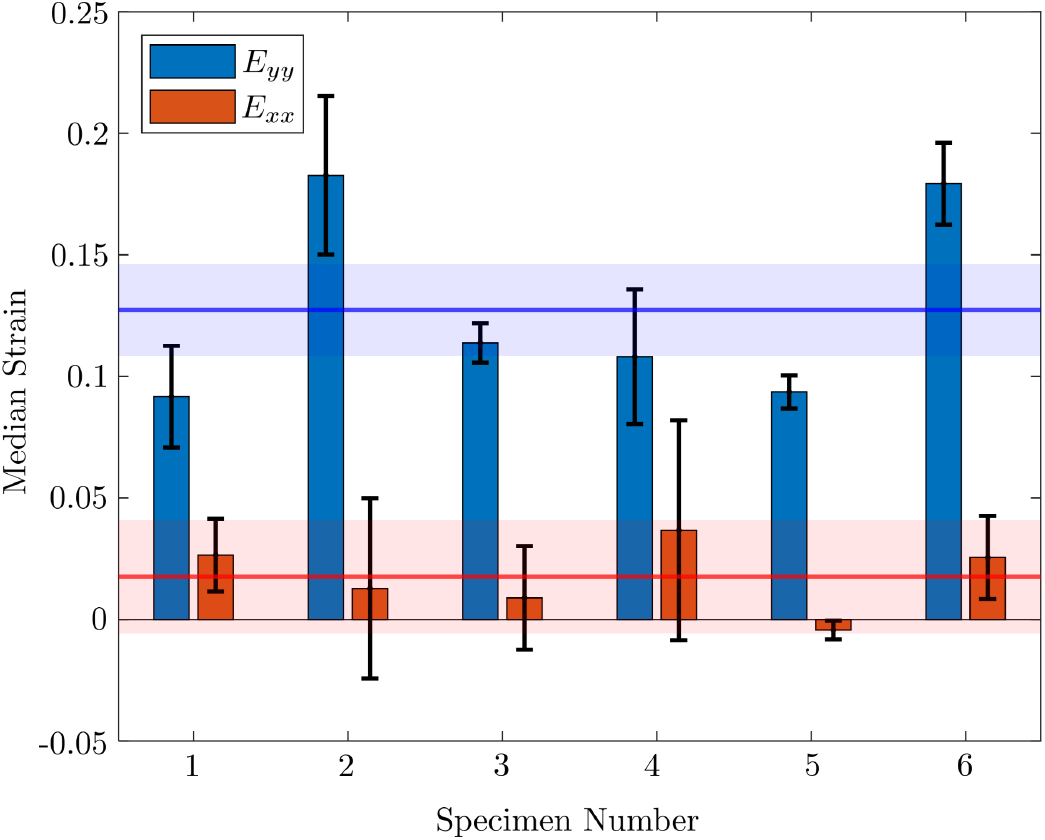
The median and interquartile range (IQR) of the Green-Lagrange strains in the circumferential *E*_*yy*_ and axial *E*_*xx*_ directions for specimens undergoing uniaxial circumferential extension, measured by Digital Image Correlation. The blue horizontal line shows the average value for median *E*_*yy*_ and the blue shaded area shows the average value for IQR of median *E*_*yy*_. The orange horizontal line shows the average value for median *E*_*xx*_ and the orange shaded area shows the average value for IQR of median *E*_*xx*_.

Additionally, we provide quantitative data within Table B.1 and Table B.2 in Appendix B. Specifically, in Table B.1 we provide the median Green-Lagrange strains and their IQRs for all six specimens undergoing DIC analysis. In Table B.2 we provide the median stretch ratios and their IQRs for all nine specimens undergoing fiber network analyses.

## 4. Discussion

We provided multiscale support to confirm the auxetic response of colorectum under uniaxial circumferential extension, a response that may provide critical insights into the underlying mechanisms of visceral pain and nociception. We utilized both DIC to optically analyze the macroscale bulk tissue deformation and our previously established fiber network analysis framework to analyze the micron-scale nerve fiber deformations during uniaxial circumferential extension of colorectal specimens [33, 34]. Both of these tools provided us valuable information and demonstrated the auxetic response of colorectal tissues.

This study provides evidence of how nature adapts to achieve favorable mechanical responses. Recent research indicates that bacteria present in the colon play a crucial role in digesting certain food components that our bodies cannot break down independently [2, 3]. This microbial digestion generates gas within the colon, which results in elevated intraluminal pressure. It is not the food content itself but rather the increased gas pressure that poses a potential risk of tissue damage [4, 5]. The auxetic response of colorectal tissues could represent an evolutionary adaptation to accommodate elevated intraluminal pressure by increasing surface area, thereby protecting the colon from excessive deformations and potential injury. This unique mechanical behavior allows the tissue to better withstand the high pressure without microstructural failure. Moreover, increased intraluminal pressure may induce further tissue distension, which closely correlates with the activation and propagation of visceral pain [7, 8]. Understanding this adaptive mechanism helps us link the mechanical response of colorectal tissue to nociception, shedding light on the evolution of protective features against stress-induced injury in the colorectum.

In the experimental setup, we kept the colorectum tubular to facilitate the delivery of relatively high-intensity displacement-driven stretching without causing tissue failure due to stress concentrations at fixation points. Conventional uniaxial or biaxial extension tests often suffer from stress concentrations at the boundaries, which limit the application of significant force or displacement [35]. We managed to deliver high-intensity circumferential displacements while maintaining the planar configuration of the myenteric plexus for successful high-resolution bright-field and fluorescent imaging.

Our experimental setup, for the first time, enabled a focused examination of circumferential and axial deformations separately, providing unique insights into the distinct mechanical behavior in each direction. Previous experimental studies either performed biaxial extension tests [20, 35, 36] or used pressure-diameter tests [37] on the tissue, which did not allow for decoupling circumferential and axial deformations.

### 4.1. Digital Image Correlation of bulk tissue deformation

We optically analyzed bulk tissue deformations during uniaxial circumferential extension, as shown in Fig. 3 and Fig. 2. The smooth stainless steel rods minimally influenced the axial load and deformation of the colorectum, allowing controlled circumferential stretch while maintaining the colorectal surface in-plane for optimal optical imaging. We observed two distinct types of tissue responses regarding axial (lateral) deformation. In five of six specimens the dominant mechanical response included significant lateral (axial) expansion during circumferential stretch. In one of six specimens the mechanical response included minimal axial (lateral) deformation during circumferential stretch.

In Fig. 2(b) we show the displacement vectors for each node derived from optical tracking, showing tissue expansion in both directions, consistent with the applied loading and auxetic response along the axial direction. The projections of the unit displacement vectors in both the *x*- and *y*-directions are substantial. The distribution of Green-Lagrange strains in Fig. 2(c),(d) demonstrate greater deformation in the *y*-direction compared to the *x*-direction which is in agreement with our expectation from mechanical loading. The distribution of axial Green-Lagrange strains (*E*_*xx*_) in Fig. 2(d) further shows positive values of *E*_*xx*_ across most of the area analyzed, providing additional evidence of the auxetic response.

In Fig. 3(b) we show the displacement vectors for each node derived from optical tracking, showing circumferential expansion consistent with the applied loading, while the projection of displacement vectors in the *x*-direction remains minimal. The distribution of Green-Lagrange strains in Fig. 3(c),(d) demonstrate greater deformation in the *y*-direction compared to the *x*-direction, emphasizing the response consistent with the loading properties and tissue unusual behavior in lateral direction.

These data provide valuable insights into the auxetic deformations of colorectal tissues, enabling a more comprehensive understanding of how the tissue responds mechanically during uniaxial extension.

### 4.2. Quantitative analyses of nerve fiber deformation

We performed live fluorescent imaging of nerve fiber networks in the colorectum under conditions of high-intensity and large-strain deformation. Our innovative use of two cylindrical rods to apply force or displacement in the circumferential direction significantly minimized movement along the through-thickness (*z*-axis), enabling high-resolution in-focus imaging of the myenteric plexus plane during deformation. We used a higher pre-stretch (relative to those using DIC) during the experiments leveraging fluorescent imaging so the signals would remain in focus, i.e. it is an expedient to avoid recording blurred fluorescent images. This design contrasts with tubular distension, where the colorectal surface continuously moves out of focus.

We utilized the VGLUT2-Cre line to drive tdTomato expression in sensory neurons, which predominantly have nerve endings concentrated in the myenteric plexus and submucosal layer [21]. The VGLUT2 promoter drives the expression of tdTomato in most colorectal sensory afferent neurons [38, 21, 39] and also a fraction (*∼*2%) of intrinsic enteric neurons [23]. We imaged the myenteric plexus from the mucosal side with high spatial and temporal resolution, successfully capturing the deformation of the complete nerve fiber network.

We implemented the resulting images into a custom algorithm to enable semi-automated analysis of the fiber network during tissue deformation, as illustrated in Fig. 5 and Fig. 4. The heatmaps of fiber stretches generated by this algorithm, in Fig. 5(b) and Fig. 4(b), allows us to visualize the distribution of micron-scale deformations across the fiber network, providing insight into their association with macroscopic, tissue-level deformations. We defined the primary range for piece-wise fiber stretch ratios (0.9-1.2) and separately highlighted the higher and lower stretch ratio ranges.

In Fig. 5(c) and Fig. 4(c) we also categorized fibers under tension and compression for separate analyses, which enhanced our understanding of their mechanical responses and provided deeper insights into the interactions within the fiber network. We then plotted the stretch ratio versus the fiber orientation relative to the axial direction (*x*-axis). In Fig. 5(d) and Fig. 4(d) we see the linear correlation between stretch ratio and orientation of unmyelinated nerve fibers shows a smaller slope in the colorectal specimen with pronounced lateral expansion compared to the specimen with almost no lateral expansion.

Our findings reveal that colorectal tissues exhibit an auxetic response, characterized by lateral expansion during circumferential stretching. This behavior likely arises from the structural organization of collagen fibers within the tissue matrix, which may inherently support auxetic deformation and enhance resistance to internal mechanical forces. Such auxetic properties have important implications for colorectal mechanotransduction, as lateral expansion may modulate how local stress and strain are transmitted to embedded sensory neurons [40]. These analyses also capture the heterogeneous deformation patterns across nerve fibers, providing key insights into the anisotropic mechanical behavior of neural tissue within the colorectum.

### 4.3. Statistical analyses

In Fig. 6 we show the primary tissue responses–median Green-Lagrange strain values– with interquartile ranges (IQR) to quantify variability across the specimens as measured using DIC. Five out of six specimens exhibited positive median *E*_*xx*_ values, indicating consistent lateral expansion during circumferential extension and confirming auxetic behavior as the dominant mechanical response in this configuration. One specimen showed a near-zero, slightly negative *E*_*xx*_, indicating minimal lateral expansion. Across all specimens, the average median *E*_*xx*_ was 0.0177, while the average median *E*_*yy*_ was 0.1273, reflecting the characteristic ratio of lateral to circumferential strain during deformation. These macroscale deformation patterns are essential for understanding the auxetic properties of the colorectum and their implications for its mechanical and physiological function.

Tables B.1 and B.2 in Appendix B present quantitative summaries of tissue and neural fiber deformation during uniaxial circumferential distension. Table B.1 reports the median Green-Lagrange strain values and corresponding interquartile ranges (IQRs) for bulk tissue, as measured using Digital Image Correlation (DIC). Table B.2 presents the median stretch ratios (*λ*) and IQRs for embedded nerve fibers, obtained through our custom fiber network analysis. These datasets collectively characterize multiscale deformations across specimens and quantify variability in mechanical response, offering key insights into both the macroscale auxetic behavior of colorectal tissue and the microscale strain experienced by embedded neural fibers.

We performed fiber network analyses on nine specimens to obtain accurate, high-resolution measurements of neural fiber deformation under circumferential stretch. Across all specimens, the average median stretch ratio was 1.0631, indicating that nerve fibers elongated by approximately 6.31% during the mechanical tests. The average interquartile range (IQR) of 0.0873 reflects substantial variability in fiber stretch, consistent with imaging data showing heterogeneous fiber orientations throughout the tissue. This variability underscores the role of fiber orientation in modulating local deformation under load. These findings offer detailed insight into the microscale mechanical behavior of neural fibers during uniaxial circumferential extension.

### 4.4. Limitations and outlook

This study has several limitations that should be acknowledged to inform future improvements. First, we conducted bulk tissue optical tracing and fiber network analyses on separate specimens. Performing both analyses on the same specimen in future studies would enable direct integration of macroscale and microscale measurements, allowing for more precise correlation between tissue-level and fiber-level responses. Second, we pre-stretched the tissue specimens to maintain the imaging focal plane during deformation. While this approach ensured consistent image quality, it may have led to a slight underestimation of stretch ratios. Nevertheless, we believe the impact on overall findings is minimal. Future studies could mitigate this limitation by employing real-time focal plane adjustment techniques, such as auto-focusing algorithms [41]. Third, we used two-dimensional image stacks of VGLUT2-labeled unmyelinated nerve fibers within the myenteric plexus, which is largely planar in structure [21, 42]. However, extending our methods to incorporate three-dimensional imaging would offer a more comprehensive understanding of tissue mechanics and fiber network morphology, including through-thickness (*z*-direction) deformation not captured in the current study [43, 44]. Addressing these limitations will enhance the ac-curacy and robustness of future investigations into colorectal tissue mechanics and nerve fiber responses.

Future directions of this work include investigating regional heterogeneity in Poisson’s ratios across the distal colorectum, encompassing colonic, intermediate, and rectal segments. Identifying spatial variations in mechanical responses may reveal structural adaptations and functional specializations within regions. Further studies using high-resolution microscale imaging could elucidate the structural origins of auxetic behavior, advancing our understanding of the mechanisms driving negative Poisson’s ratios in biological tissues. In addition, changes in Poisson’s ratio may serve as a novel biomechanical marker for tissue health, offering a potential diagnostic tool for detecting pathological alterations in colorectal mechanics [17]. The foundational data presented here support continued experimental and computational studies aimed at uncovering how biological systems achieve auxeticity. Insights gained from these mechanisms could inform the design of bioinspired auxetic materials and mechanical metamaterials with enhanced functionality [45]. Such advances hold promise for transformative applications in engineering and biomedicine, where materials mimicking the exceptional properties of biological tissues are increasingly in demand.

## Acknowledgment

NSF 1727185 and NIH 1R01DK120824-01.

During the preparation of this work the authors used ChatGPT-4 to sparingly edit for brevity. The authors reviewed and edited the content in detail after using this tool/service and take full responsibility for the content of the publication.

## Appendix A. Supplemental images of nerve fiber deformation

In Fig. A.7 we show a representative image of fluorescent nerve fibers from a VG-LUT2/tdT mouse colorectum in the initial and final configurations.

**Figure A7:**
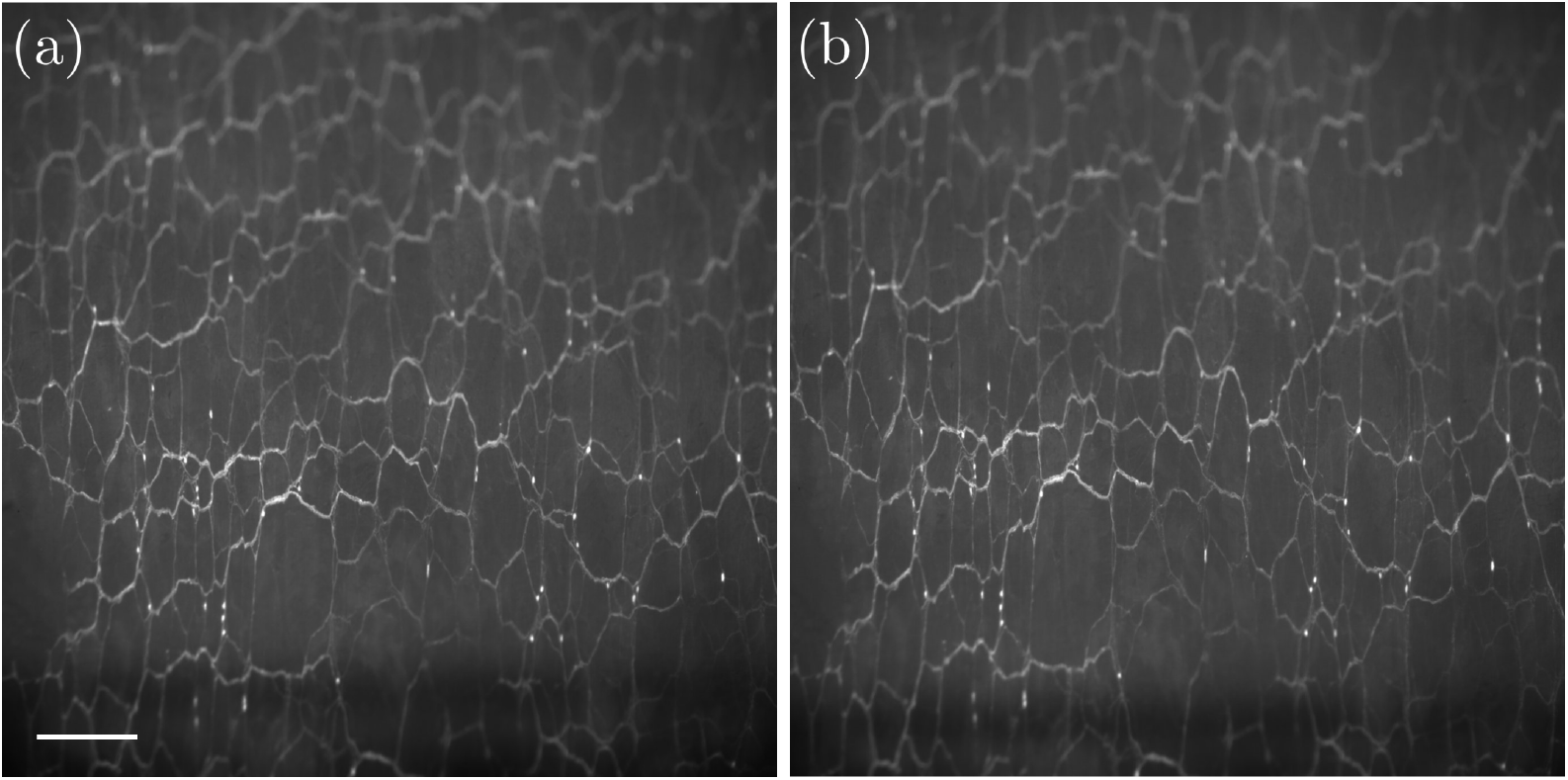
Representative magnified view of fluorescent microscopy images of VGLUT2-labeled nerve fibers during circumferential extension of the distal colon. (a) Initial configuration, and (b) corresponding final configuration at maximum displacement. The scale bar equals 465 *µ*m.

## Appendix B. Supplemental quantitative analyses

In the following tables, we present the results of statistical analyses performed on our cohort of specimens after conducting analyses leveraging DIC (Table B.1) and leveraging our custom fiber network analyses (Table B.2).

**Table B.1:** The median strain and interquartile range (IQR) for specimens undergoing uniaxial circumferential extension and DIC analyses.

| Specimen | Strain | Median | IQR |
| --- | --- | --- | --- |
| 1 | $E_{yy}$ | 0.0917 | 0.0209 |
| | $E_{xx}$ | 0.0265 | 0.0149 |
| 2 | $E_{yy}$ | 0.1827 | 0.0326 |
| | $E_{xx}$ | 0.0128 | 0.0371 |
| 3 | $E_{yy}$ | 0.1138 | 0.0081 |
| | $E_{xx}$ | 0.0089 | 0.0213 |
| 4 | $E_{yy}$ | 0.1081 | 0.0277 |
| | $E_{xx}$ | 0.0367 | 0.0452 |
| 5 | $E_{yy}$ | 0.0936 | 0.0068 |
| | $E_{xx}$ | -0.0043 | 0.0039 |
| 6 | $E_{yy}$ | 0.1793 | 0.0169 |
| | $E_{xx}$ | 0.0255 | 0.0171 |
| **Average** | $E_{yy}$ | 0.1273 | 0.0188 |
| | $E_{xx}$ | 0.0177 | 0.0233 |

**Table B.2:** The median stretch ratio (*λ*) and interquartile range (IQR) for specimens undergoing uniaxial circumferential extension and fiber network analyses.

| Specimen | Stretch ratio (median) | Stretch ratio (IQR) |
| --- | --- | --- |
| 1 | 1.0674 | 0.0729 |
| 2 | 1.0892 | 0.0784 |
| 3 | 1.0661 | 0.0938 |
| 4 | 1.0788 | 0.0978 |
| 5 | 1.0444 | 0.0840 |
| 6 | 1.0497 | 0.0914 |
| 7 | 1.0519 | 0.0969 |
| 8 | 1.0660 | 0.0797 |
| 9 | 1.0543 | 0.0911 |
| **Average** | **1.0631** | **0.0873** |

## References

[1] E. Scarpellini, L. M. Balsiger, B. Broeders, K. V. D. Houte, K. Routhiaux, K. Raymenants, F. Carbone, J. Tack, Nutrition and disorders of gut–brain interaction, Nutrients 16 (1) (2024) 176.

[2] G. Barbara, C. Feinle-Bisset, U. C. Ghoshal, J. Santos, S. J. Vanner, N. Vergnolle, E. G. Zoetendal, E. M. Quigley, The intestinal microenvironment and functional gastrointestinal disorders, Gastroenterology 150 (6) (2016) 1305–1318.

[3] C. R. Singh, H. Zogg, S. Ro, Role of micrornas in disorders of gut–brain interactions: clinical insights and therapeutic alternatives, Journal of Personalized Medicine 11 (10) (2021) 1021.

[4] D. R. A. Mars, M. Frith, P. C. Kashyap, Functional gastrointestinal disorders and the microbiome—what is the best strategy for moving microbiome-based therapies for functional gastrointestinal disorders into the clinic?, Gastroenterology 160 (2) (2021) 538–555.

[5] L. Kader, A. B. Willits, S. Meriano, J. A. Christianson, J.-H. La, B. Feng, B. Knight, G. Kosova, J. J. Deberry, M. D. Coates, et al., Identification of arginine-vasopressin receptor 1a (avpr1a/avpr1a) as a novel candidate gene for chronic visceral pain sheds light on the potential role of enteric neurons in the development of visceral hypersensitivity, The Journal of Pain 25 (9) (2024) 104572.

[6] P. J. Pasricha, W. D. Willis, G. F. Gebhart, Chronic abdominal and visceral pain: theory and practice, CRC Press, 2006.

[7] A. Agrawal, P. Whorwell, abdominal bloating and distension in functional gastrointestinal disorders–epidemiology and exploration of possible mechanisms, Alimentary pharmacology & therapeutics 27 (1) (2008) 2–10.

[8] Q. Zhou, G. N. Verne, New insights into visceral hypersensitivity—clinical implications in ibs, Nature Reviews Gastroenterology & Hepatology 8 (6) (2011) 349–355.

[9] B. Feng, P. R. Brumovsky, G. F. Gebhart, Differential roles of stretch-sensitive pelvic nerve afferents innervating mouse distal colon and rectum, American Journal of Physiology-Gastrointestinal and Liver Physiology 298 (3) (2010) G402–G409.

[10] B. Feng, T. Guo, Visceral pain from colon and rectum: the mechanotransduction and biomechanics, Journal of Neural Transmission 127 (4) (2020) 415–429.

[11] A. C. Ford, S. Vanner, P. C. Kashyap, Y. Nasser, Chronic visceral pain: new peripheral mechanistic insights and resulting treatments, Gastroenterology 166 (6) (2024) 976– 994.

[12] A. Shokrani, A. Almasi, B. Feng, D. M. Pierce, Understanding mechanotransduction in the distal colon and rectum via multiscale and multimodal computational modeling, Journal of the Mechanical Behavior of Biomedical Materials (2024) 106771.

[13] S. Adeeb, A. Ali, N. Shrive, C. Frank, D. Smith, Modelling the behaviour of ligaments: a technical note, Computer methods in biomechanics and biomedical engineering 7 (1) (2004) 33–42.

[14] E. Danso, P. Julkunen, R. Korhonen, Poisson’s ratio of bovine meniscus determined combining unconfined and confined compression, Journal of biomechanics 77 (2018) 233–237.

[15] P. Skacel, J. Bursa, Poisson’s ratio and compressibility of arterial wall–improved experimental data reject auxetic behaviour, Journal of the Mechanical Behavior of Biomedical Materials 131 (2022) 105229.

[16] C. Lees, J. F. Vincent, J. E. Hillerton, Poisson’s ratio in skin, Bio-medical materials and engineering 1 (1) (1991) 19–23.

[17] R. Gatt, M. V. Wood, A. Gatt, F. Zarb, C. Formosa, K. M. Azzopardi, A. Casha, T. P. Agius, P. Schembri-Wismayer, L. Attard, et al., Negative poisson’s ratios in tendons: An unexpected mechanical response, Acta biomaterialia 24 (2015) 201–208.

[18] N. J. Spencer, M. Kyloh, M. Duffield, Identification of different types of spinal afferent nerve endings that encode noxious and innocuous stimuli in the large intestine using a novel anterograde tracing technique, PloS one 9 (11) (2014) e112466.

[19] S. Brookes, N. Chen, A. Humenick, N. J. Spencer, M. Costa, Extrinsic sensory innervation of the gut: structure and function, The Enteric Nervous System: 30 Years Later (2016) 63–69.

[20] S. Siri, F. Maier, S. Santos, D. M. Pierce, B. Feng, Load-bearing function of the colorectal submucosa and its relevance to visceral nociception elicited by mechanical stretch, American Journal of Physiology-Gastrointestinal and Liver Physiology 317 (3) (2019) G349–G358.

[21] T. Guo, S. Patel, D. Shah, L. Chi, S. Emadi, D. M. Pierce, M. Han, P. R. Brumovsky, B. Feng, Optical clearing reveals tnbs-induced morphological changes of vglut2-positive nerve fibers in mouse colorectum, American Journal of Physiology-Gastrointestinal and Liver Physiology 320 (4) (2021) G644–G657.

[22] B. Feng, L. Chen, S. J. Ilham, A review on ultrasonic neuromodulation of the peripheral nervous system: enhanced or suppressed activities?, Applied Sciences 9 (8) (2019) 1637.

[23] J. Liu, S. Zhang, S. Emadi, T. Guo, L. Chen, B. Feng, Morphological, molecular, and functional characterization of mouse glutamatergic myenteric neurons, American Journal of Physiology-Gastrointestinal and Liver Physiology 326 (3) (2024) G279–G290.

[24] B. Feng, J. H. La, E. S. Schwartz, G. F. Gebhart, Irritable bowel syndrome: methods, mechanisms, and pathophysiology. neural and neuro-immune mechanisms of visceral hypersensitivity in irritable bowel syndrome, American Journal of Physiology-Gastrointestinal and Liver Physiology 302 (10) (2012) G1085–G1098.

[25] Y. Zhao, S. Siri, B. Feng, D. M. Pierce, The macro-and micro-mechanics of the colon and rectum II: Theoretical and computational methods, Bioengineering 7 (4) (2020) 152.

[26] A. Shokrani, A. Almasi, B. Feng, D. M. Pierce, Corrigendum to “predicting the micromechanics of embedded nerve fibers using a novel three-layered model of mouse distal colon and rectum” [J. Mech. Beha. Biomed. Mater. 127 (2022) 105083], Journal of the mechanical behavior of biomedical materials 154 (2024) 106286.

[27] F. Cervero, J. M. Laird, Visceral pain, The Lancet 353 (9170) (1999) 2145–2148.

[28] H. Eilers, M. A. Schumacher, Mechanosensitivity of primary afferent nociceptors in the pain pathway, in: A. Kamkin, I. Kiseleva (Eds.), Mechanosensitivity in Cells and Tissues, Academia, 2005.

[29] B. Feng, J.-H. La, T. Tanaka, E. S. Schwartz, T. P. McMurray, G. F. Gebhart, Altered colorectal afferent function associated with tnbs-induced visceral hypersensitivity in mice, American Journal of Physiology-Gastrointestinal and Liver Physiology 303 (7) (2012) G817–G824.

[30] A. D. Edelstein, M. A. Tsuchida, N. Amodaj, H. Pinkard, R. D. Vale, N. Stuurman, Advanced methods of microscope control using μmanager software, Journal of biological methods 1 (2) (2014) e10.

[31] J. Humphrey, S. DeLange, An Introduction to Biomechanics: Solids and Fluids, Analysis and Design, Springer Link, 2015.

[32] A. Shokrani, A. Seck, B. Feng, D. Pierce, Methods for quantitative analyses of nerve fiber deformation in the myenteric plexus under loading of mouse distal colon and rectum (submitted for review).

[33] D. Jaiswal, N. Cowley, Z. Bian, G. Zheng, K. P. Claffey, K. Hoshino, Stiffness analysis of 3d spheroids using microtweezers, PloS one 12 (11) (2017) e0188346.

[34] D. Jaiswal, M. D. Tang-Schomer, D. Sood, D. L. Kaplan, K. Hoshino, Nondestructive, label-free characterization of mechanical microheterogeneity in biomimetic materials, ACS Biomaterials Science & Engineering 4 (9) (2018) 3259–3267.

[35] S. Siri, F. Maier, L. Chen, S. Santos, D. M. Pierce, B. Feng, Differential biomechanical properties of mouse distal colon and rectum innervated by the splanchnic and pelvic afferents, American Journal of Physiology-Gastrointestinal and Liver Physiology 316 (4) (2019) G473–G481.

[36] S. Puértolas, E. Peña, A. Herrera, E. Ibarz, L. Gracia, A comparative study of hyperelastic constitutive models for colonic tissue fitted to multiaxial experimental testing, Journal of the mechanical behavior of biomedical materials 102 (2020) 103507.

[37] D. P. Sokolis, S. G. Sassani, Microstructure-based constitutive modeling for the large intestine validated by histological observations, Journal of the mechanical behavior of biomedical materials 21 (2013) 149–166.

[38] T. Guo, Z. Bian, K. Trocki, L. Chen, G. Zheng, B. Feng, Optical recording reveals topological distribution of functionally classified colorectal afferent neurons in intact lumbosacral drg, Physiological reports 7 (9) (2019) e14097.

[39] T. Guo, J. Liu, L. Chen, Z. Bian, G. Zheng, B. Feng, Sex differences in zymosan-induced behavioral visceral hypersensitivity and colorectal afferent sensitization, American Journal of Physiology-Gastrointestinal and Liver Physiology 326 (2) (2024) G133–G146.

[40] B. Feng, Y. Zhu, J.-H. La, Z. P. Wills, G. F. Gebhart, Experimental and computational evidence for an essential role of nav1. 6 in spike initiation at stretch-sensitive colorectal afferent endings, Journal of neurophysiology 113 (7) (2015) 2618–2634.

[41] Y. Sun, S. Duthaler, B. J. Nelson, Autofocusing algorithm selection in computer microscopy, in: 2005 IEEE/RSJ International Conference on Intelligent Robots and Systems, IEEE, 2005, pp. 70–76.

[42] T. Guo, S. Patel, D. Shah, L. Chi, S. Emadi, D. M. Pierce, M. Han, P. R. Brumovsky, B. Feng, Neurogastroenterology and motility: Optical clearing reveals tnbs-induced morphological changes of vglut2-positive nerve fibers in mouse colorectum, American Journal of Physiology-Gastrointestinal and Liver Physiology 320 (4) (2021) G644.

[43] Y.-A. Liu, Y.-C. Chung, S.-T. Pan, Y.-C. Hou, S.-J. Peng, P. J. Pasricha, S.-C. Tang, 3-d illustration of network orientations of interstitial cells of cajal subgroups in human colon as revealed by deep-tissue imaging with optical clearing, American Journal of Physiology-Gastrointestinal and Liver Physiology 302 (10) (2012) G1099–G1110.

[44] Y. Liu, Y. Chung, S. Pan, M. Shen, Y. Hou, S. Peng, P. Pasricha, S. Tang, 3-d imaging, illustration, and quantitation of enteric glial network in transparent human colon mucosa, Neurogastroenterology & Motility 25 (5) (2013) e324–e338.

[45] J. N. Grima, R. Caruana-Gauci, Materials that push back, Nature materials 11 (7) (2012) 565–566.

